# Effective cell membrane tension is independent of polyacrylamide substrate stiffness

**DOI:** 10.1101/2021.11.09.467973

**Authors:** Eva Kreysing, Jeffrey Mc Hugh, Sarah K. Foster, Kurt Andresen, Ryan D. Greenhalgh, Eva K. Pillai, Andrea Dimitracopoulos, Ulrich F. Keyser, Kristian Franze

## Abstract

Most animal cells are surrounded by a cell membrane and an underlying actomyosin cortex. Both structures are linked, and they are under tension. In-plane membrane tension and cortical tension both influence many cellular processes, including cell migration, division, and endocytosis. However, while actomyosin tension is regulated by substrate stiffness, how membrane tension responds to mechanical substrate properties is currently poorly understood. Here, we probed the effective membrane tension of neurons and fibroblasts cultured on glass and polyacrylamide substrates of varying stiffness using optical tweezers. In contrast to actomyosin-based traction forces, both peak forces and steady state tether forces of cells cultured on hydrogels were independent of substrate stiffness and did not change after blocking myosin II activity using blebbistatin, indicating that tether and traction forces are not directly linked. Peak forces in fibroblasts on hydrogels were about twice as high as those in neurons, indicating stronger membrane-cortex adhesion in fibroblasts. Steady state tether forces were generally higher in cells cultured on hydrogels than on glass, which we explain by a mechanical model. Our results provide new insights into the complex regulation of effective membrane tension and pave the way for a deeper understanding of the biological processes it instructs.

## Introduction

The cell membrane is supported by the underlying actomyosin cortex, and molecular linkers form tight connections between the two layers. Osmotic pressure resulting from ion gradients across the membrane and interactions of the membrane with the force-generating actomyosin cytoskeleton contribute to a tension across the membrane. This membrane tension regulates many cellular processes, such as cell migration, division, stem cell fate choice, and endo- and exocytosis^1–6^. Despite its biological importance^7–9^, membrane tension is still difficult to quantify. In-plane membrane tension is defined as the force per unit length acting tangentially on the plasma membrane. Because of the membrane’s coupling to cytoskeletal elements, measurements of the ‘effective’ membrane tension usually contain contributions of the underlying actomyosin cortex and other force-generating elements^10,11^.

The actomyosin cortex is coupled to the extracellular environment through transmembrane proteins such as integrins and cadherins, which transmit tensile forces generated by acto-myosin interaction to the environment. The resulting traction forces, and hence cytoskeletal tension, increase with substrate stiffness for most cell types^12,13^, suggesting that the measured effective membrane tension should also increase on stiffer substrates. However, the regulation of membrane tension by substrate mechanics is yet to be probed experimentally.

## Results

Here, we used optical tweezers (OT) to probe membrane properties of cells cultured on polyacrylamide substrates of different stiffnesses as well as on glass. Using a membrane-adherent bead held in an optical trap, we pulled membrane tethers from cells, and the resulting tether forces served as an indicator for effective membrane tension ^2,14,15^ (Fig. 1a, b; Fig. S1, Video S1). Throughout a pull, the force on the tether can be continuously monitored, resulting in a characteristic force-time curve (Fig. 1c). The force rises quickly after initiation of the pull and reaches a peak before decreasing sharply. In agreement with Sheetz^14^, we attribute this peak force (PF) to the initial local detachment of plasma membrane from the actomyosin cortex (Fig. 1a), although it may also be influenced by measurement parameters such as the adhesion area between the bead and the cell membrane^16^.

**Figure 1:**
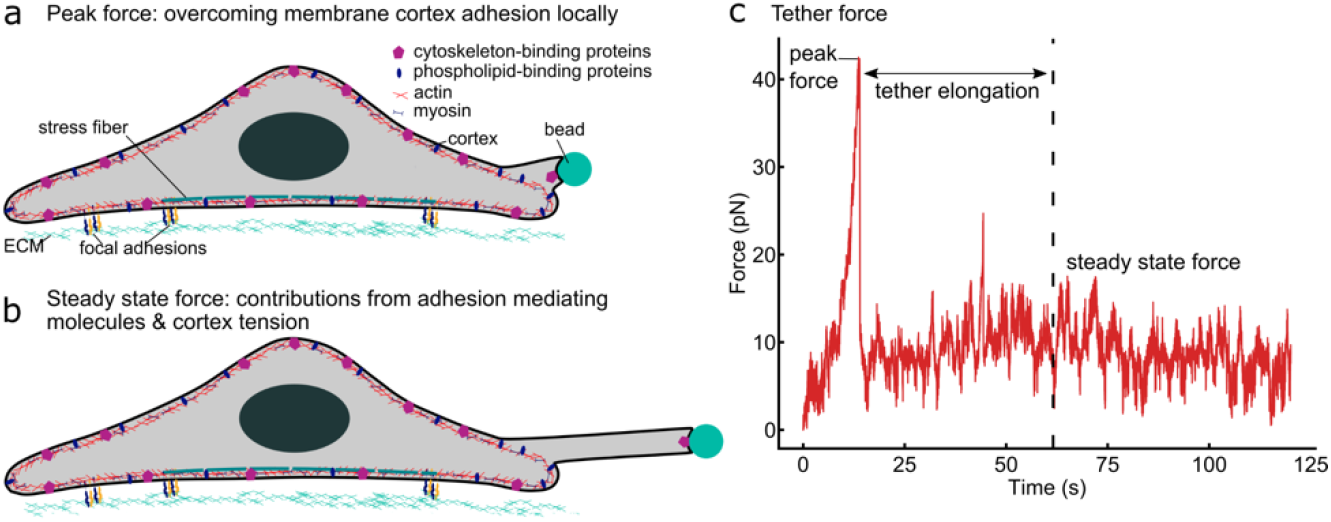
Illustration of an OT measurement. Peak and steady state tether forces provide different insights into cell membrane properties. (a) When an OT pull is initiated, the local membrane-cortex adhesion first needs to be overcome, resulting in a peak force (PF) measured at the beginning of the experiment (see (c)). (b) After tether extension, the bead is held stationary, and the steady state force (SSF), which scales with effective membrane tension, is recorded. (c) A typical force-time curve recorded during a tether pull, showing a peak (PF) at the beginning of the pull and a plateau (SSF) when the tether is held at its maximum extension.

Subsequently, the tether is pulled until it reaches a set length. During this elongation process, the plasma membrane slides over the actin cortex while bonds between the membrane and the lipid-binding proteins located in or at the cortex are broken and quickly re-established^17^. At the same time, membrane proteins may accumulate at the base of the tether and influence the pulling force^15^. Overall, the measured forces then decrease during the elongation phase. Afterwards, the bead is held in a stable position and the steady-state force (SSF) is measured (Fig. 1b). The SSF is thought to reflect a combination of the in-plane plasma membrane tension and the cortical tension, with the strength of the membrane-cortex adhesion determining the coupling between these two elements^17,18^. As such, the SSF depends in a non-linear manner on the number of cortex-plasma membrane linkers such a ERM proteins^19^.

To test the effect of substrate mechanics and cortical tension on effective membrane tension, we performed tether pulls in fibroblasts, which possess a dense actomyosin cortex, and in neuronal axons, whose sub-membranous cytoskeleton is characterized by actin rings spaced by spectrin tetramers^20^. Cells were cultured on glass as well as on hydrogels fabricated within a stiffness range adjusted to match their natural environment. For fibroblasts, hydrogel stiffnesses ranged from 100Pa (very soft; similar to subcutaneous adipose tissue), to 10kPa (stiff; already in the stiffness range of many fibrotic tissues)^21^. For neurons, the highest substrate stiffness used was 1kPa, corresponding to the highest stiffness experienced by this type of neuron *in vivo^22^*.

Peak forces (PFs) measured in fibroblasts were about twice as high as those measured in neuronal axons (Fig. 2b, f). Within each cell type, PFs were similar across all tested hydrogel substrates. Steady state forces (SSFs), which were comparable between fibroblasts and neurons, were also similar across all polyacrylamide substrate stiffnesses (Fig. 2c, g).

**Figure 2:**
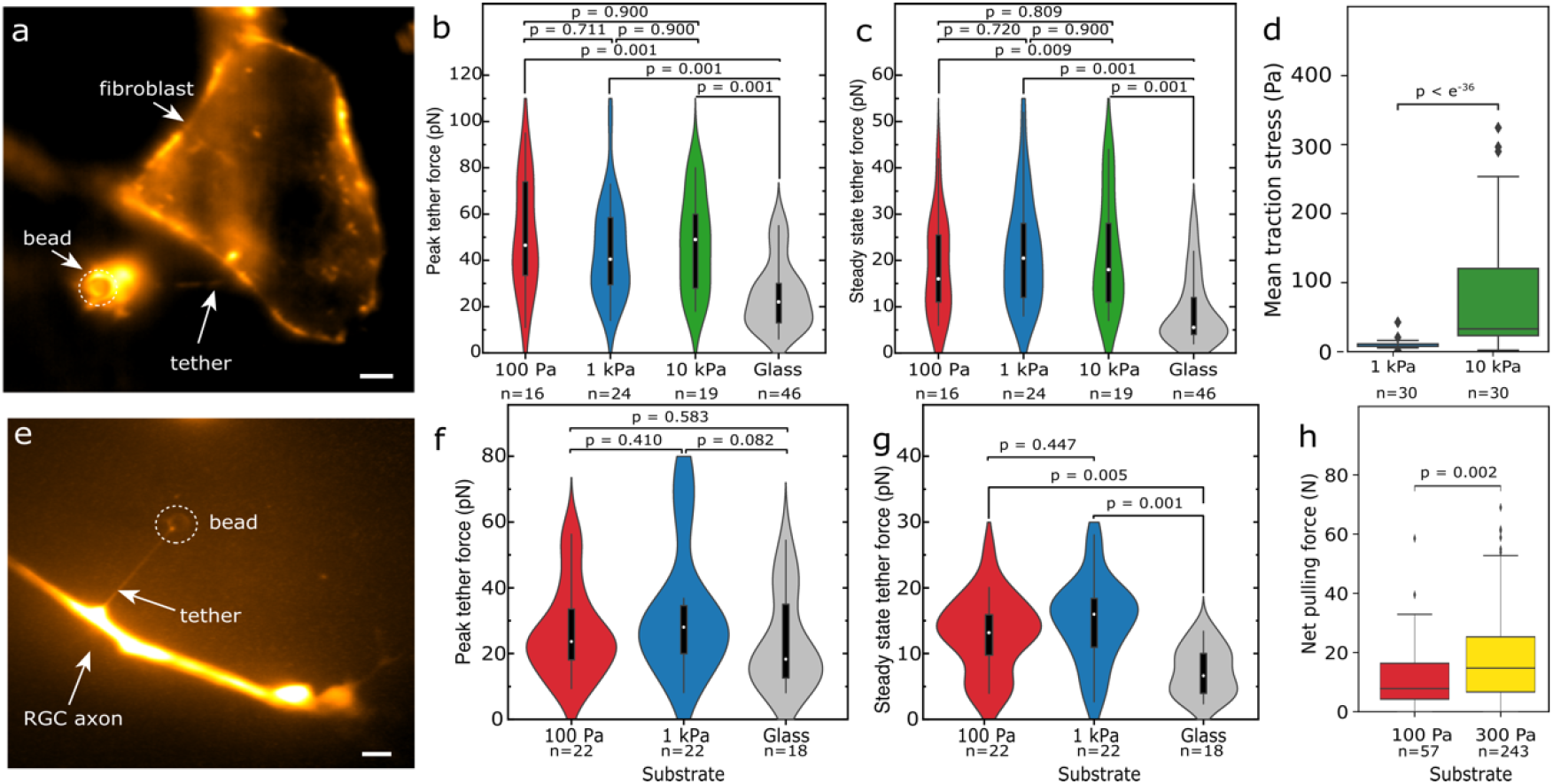
Tether and traction forces of fibroblasts and neurons on different substrates. (a-d) 3T3 fibroblasts and (e-h) *Xenopus* retinal ganglion cell axons; shear moduli of polyacrylamide substrates are provided in Pa. (a, e) Fluorescence images of membrane tethers pulled from a 3T3 fibroblast and an axon. Fibroblast membranes were labelled with CellMask membrane stain, neurons with Tetramethylrhodamine Dextran. Scale bars: 5μm. (b) Peak tether forces (PFs) and (c) steady state tether forces (SSFs) were similar on all hydrogels irrespective of their stiffness (Kruskal-Wallis tests followed by Tukey posthoc tests), but significantly larger than on glass. (d) 3T3 fibroblasts exerted higher traction forces on stiffer hydrogels compared to on softer ones (two-tailed Mann-Whitney test). (f) PFs in axons were similar on all hydrogel and glass substrates. (g) SSFs in neurons were similar on different hydrogels but higher than those measured on glass. (h) Net forces (growth cone forces pulling on the axon) were higher on stiffer hydrogels than on softer ones (two-tailed Mann-Whitney test).

In contrast, actomyosin-based traction forces, i.e., contractile cellular forces exerted on the substrate, are known to scale with substrate stiffnesses for both neurons and fibroblasts^23,24^. We measured these forces in our culture conditions using traction force microscopy (TFM), where cells are cultured on soft, elastic hydrogels with embedded fluorescent nanoparticles (see Methods)^25^. Forces exerted by the cells lead to local substrate deformations, which can be tracked using fluorescence imaging of the nanoparticles, whose displacement can be used to calculate traction forces^26^.

We confirmed that traction forces exhibited by both cell types increased significantly with increasing substrate stiffness within the investigated stiffness range (Fig. 2d, Fig. S2). Similarly, net forces, which are obtained by integrating traction forces across neuronal growth cones, and which provide an approximation for the tension exerted by a growth cone on its axon^23,27^, significantly increased on stiffer substrates (Fig 2h). These data indicated that traction forces and net forces assessed by 2D traction force microscopy and tether forces measured by OT are not tightly coupled.

In order to further examine the interplay between cortical tension and effective membrane tension, we measured tether forces in fibroblasts cultured on 10 kPa hydrogels, where we would expect the highest cytoskeletal tension^24^, following treatment with the myosin II (and thus contractility) inhibitor blebbistatin. Blebbistatin treatment had no significant effect on either PFs or SSFs (Fig. 3), suggesting that there is indeed no strong coupling between actomyosin-based cortical tension and average effective membrane tension in cells cultured on hydrogels.

**Figure 3:**
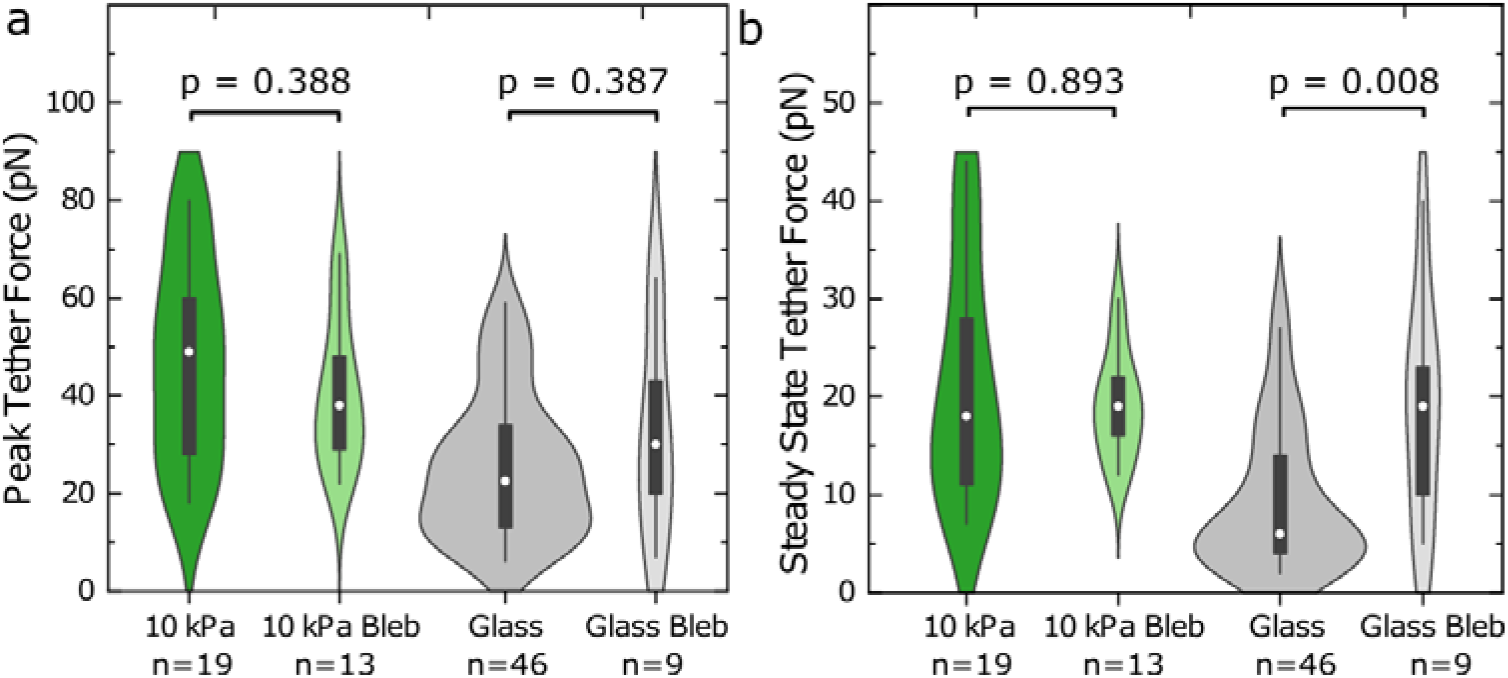
Inhibition of actomyosin contractility increases SSFs on glass but not on hydrogels. (a) The application of 20 μM Blebbistatin did not significantly alter PFs on hydrogels or on glass (Mann-Whitney test). (b) SSFs of fibroblasts grown on glass but not on hydrogels increased significantly following blebbistatin treatment. The untreated conditions correspond to the data shown in Figure 2.

As numerous *in vitro* studies use glass-bottomed culture dishes, and the surface properties of glass differ considerably from those of hydrogels, we also cultured fibroblasts and neurons on glass to test whether effective membrane tension is comparable in both culture environments.

In neurons, PFs measured in axons cultured on glass were indeed like those observed on hydrogels (Fig. 2f). However, PFs for fibroblasts cultured on glass were significantly lower than those for fibroblasts grown on hydrogels (Fig. 2b). Similarly, SSFs were significantly lower for fibroblasts and neurons cultured on glass compared to those cultured on hydrogels (Fig. 2c, g). Unlike on hydrogels, tether forces measured on glass were similar for fibroblasts and neurons (Fig. 2).

While PFs did not change significantly after blocking actomyosin contractility in fibroblasts cultured on glass, SSFs increased on glass following blebbistatin treatment^28^ (Fig 3b), reaching values similar to SSFs seen in fibroblasts cultured on hydrogels. These data showed that effective membrane tension depends on the type of culture substrate, and they suggested that the coupling between the cell cortex and the membrane also differs between glass and hydrogel substrates.

## Discussion

Here we found that the steady-state tether forces, which are thought to scale with effective membrane tension, do not change as a function of hydrogel substrate stiffness within a physiologically relevant stiffness range in both fibroblasts and neurons (Fig. 2c, g). In contrast, actomyosin-based traction and net forces increased on stiffer substrates (Fig. 2d, h), as shown previously*^24^*, suggesting that the effective membrane tension measured by OT dorsolaterally to adhesion sites is not directly linked to cellular contractility on hydrogels*^29^*. In line with this interpretation, the application of the myosin II blocker, blebbistatin, did not alter the effective membrane tension of fibroblasts cultured on hydrogels (Fig. 3).

Alternatively, membrane tension might be influenced by cortical tension on short time scales but is then actively regulated to resume original values quickly*^30^*. Another possibility is that traction forces and membrane tension are coupled also on longer time scales but traction forces transmitted from focal adhesions and ventral stress fibres to the actin cortex*^31^* dissipate with increasing distance from the stress fibres. Thus, on hydrogels enhanced cortical tension might be limited to ventral areas of the cell surface not accessible by OT. This would be in line with a recent study suggesting that cortical tension in dorsolateral parts of the cell are largely independent of traction forces^24^.

If cortical and membrane tension are coupled, membrane tension would then only change locally^32^ near focal adhesions and stress fibres^31^, which would not be seen when probed further away. Such local membrane tension heterogeneities would explain, for example, why the mechanosensitive ion channel Piezo1, which is thought to be activated by membrane tension, is activated only near focal adhesion sites despite being distributed across the whole cell surface^33^.

The effective membrane tension was higher in cells cultured on hydrogels than in those cultured on glass (Fig. 2). Fibroblasts developed much stronger stress fibres on glass than on hydrogels (Fig. S3)*^34^*, indicating considerably larger traction forces. The strong contraction of stress fibres and the embedding actin cortex*^33^* on glass could propagate further within the actin cortex than on hydrogels. The concomitant decrease in cortex area could cause the membrane to relax also in areas assessed by OT in cells cultured on glass, as schematically indicated in Figure 4, explaining why tether forces were lower on glass than on hydrogels (Fig. 2).

**Figure 4:**
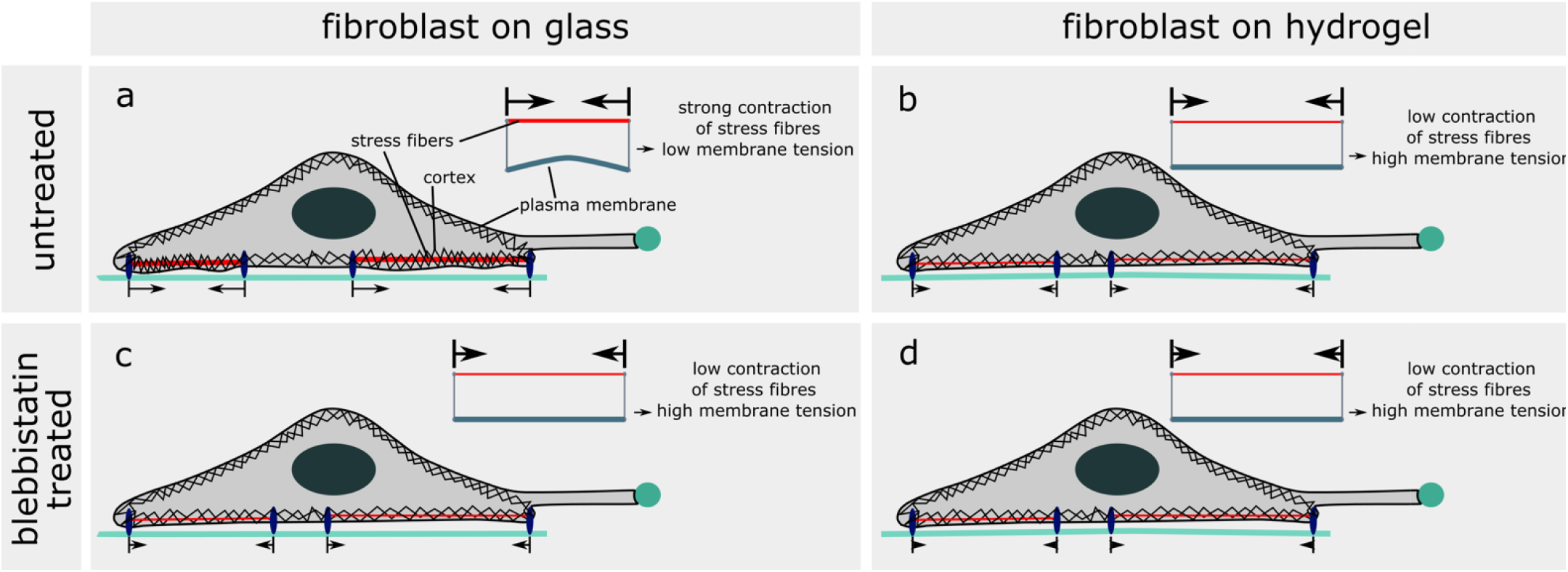
Toy model of substrate type-dependent effective membrane tension. Schematic representation of the cell cortex, stress fibres and cell membrane in fibroblasts on glass and hydrogel substrates without and with blebbistatin treatment. (a) In fibroblasts on glass substrates without blebbistatin treatment, the strong contraction of stress fibres, which is transmitted to the nearby actin cortex^31^, might lead to the relaxation of the membrane and thus reduce membrane tension. (b) In fibroblasts on hydrogel substrates without blebbistatin treatment, we observed fewer stress fibres, most likely resulting in lower contractility and less relaxation of the membrane. The resulting membrane tension is higher compared to that of cells on glass. (c) In fibroblasts on glass substrates with blebbistatin treatment, the stress fibres relax. This might increase membrane tension. (d) In fibroblasts on hydrogel substrates with blebbistatin treatment, fewer stress fibres relax, and the effect on membrane tension is too low to be detected by OT.

This could not only explain why SSFs in fibroblasts were lower on glass than on hydrogels (Fig. 2c, Fig. 4b) but also why blebbistatin treatment, which relaxes both stress fibres, led to a significant increase in SSFs on glass but not on hydrogels (Figs. 3b, 4c-d). The tension exerted by growth cones on the distal axon, which increased on stiffer substrates (Fig. 2h), could similarly lead to membrane relaxation in neurons cultured on glass (Fig. 2g). Alternatively, or additionally, differences in physical and chemical properties between glass and hydrogels could also activate different signalling pathways involved in the regulation of membrane tension homeostasis^30^.

The PF, which is the initial force required to detach the membrane from the cortex, is related to membrane-cortex adhesion^14^ (in analogy to attempting to open Velcro from the middle). Fibroblasts have a dense actomyosin cortex, and proteins such as ezrin/radixin/moesin (ERM proteins) can link the membrane to the cortex across the cell surface. Neuronal axons, on the other hand, have a periodic actin-spectrin network underneath their membrane^20^ and much less area for ERM proteins to connect the membrane to actin rings. This morphological difference may lead to lower membrane-cortex adhesion in axons compared to fibroblasts, explaining why PFs are about twice as high in fibroblasts compared to axons cultured on physiologically stiff hydrogels. A low membrane-cortex coupling could also explain why tethers pulled from axons can freely “slide” along the axon as observed in a previous and the current study (video S2). To our knowledge, tether sliding has not been reported in fibroblasts.

Our results confirm that membrane tension measurements need to be interpreted carefully. Effective membrane tension dorsolaterally of adhesion sites did not depend on substrate stiffness within a physiological stiffness range, yet it varied between hydrogels and glass, and between cell types. Future technological developments enabling highly resolved quantitative membrane tension measurements will shed light on the molecular control of local membrane tension, an important regulator of many cellular functions.

## Acknowledgments

The authors would like to thank Alex Winkel and Joy Thompson for AFM measurements of hydrogels, Liz Williams for preparation of hydrogels, and Ewa Paluch, Ruby Peters and Aki Stubb for discussions. J. Mc H. acknowledges funding from AFOSR (Grant No. FA9550-17-1-0118). S. K. F. acknowledges funding from the Herchel Smith Foundation. U. F. K. was supported by an ERC Consolidator Grant DesignerPores No. 647144, K. F. was supported by the European Research Council (Consolidator Grant 772426) and the Alexander von Humboldt Foundation (Alexander von Humboldt Professorship).

## Author Contributions

J. Mc H., E. K., and K. A. performed tether pulling experiments. J. Mc H. analysed the results from OT experiments. E. K. cultured 3T3 fibroblasts and prepared hydrogels. S. K. F. dissected and cultured Xenopus laevis RGCs and prepared hydrogels. A.D. performed TFM experiments with 3T3 fibroblasts. R. G. performed the TFM experiments with neurons and all TFM analyses. E. K. and E. P. performed the immunostainings. J. Mc H., E. K., K. F., and U. F. K. designed experiments with 3T3 fibroblasts. J. Mc H., S. K. F., K. A., and U. F. K. designed pulling experiments with X. laevis RGCs. S. K. F., J. Mc H., E. K., U.F.K. and K. F. conceived the project. E. K., J. Mc H., S. K. F., U.F.K. and K.F. wrote the manuscript.

## Materials and methods

### NIH 3T3 fibroblast cell culture

3T3 cells were cultured in plastic cell culture flasks in culture media (90 ml DMEM with glutamine (Gibco 22320-022), 10 ml Fetal Bovine Serum (Gibco 25300054), 1 ml Penicillin-Streptomycin-Amphotericin B Mixture (PSF, Lonza 17-745E)). After reaching confluency, cells were enzymatically detached. For this step, cells were washed twice with prewarmed (37°C) PBS (Sigma D8537) and then incubated with 1ml 10% TrypLE (Thermo Fisher Scientific A1285901) in PBS at 37°C for 5 min. 5 ml culture media was then added, and the cells were cultured in fresh culture flasks or plated on glass dishes and polyacrylamide (PAA) hydrogels for 4-48 h prior to experiments.

Hydrogels of 3 different elastic moduli (shear moduli = 100 Pa,1 kPa, and 10 kPa) were prepared and functionalized as described below. PAA gels as well as glass bottom culture dishes were coated with 0.01 mg/ml PDL in PBS for at least 4 h (typically overnight) followed by 0.02 mg/ml fibronectin in PBS for at least 4 h. Culture media was exchanged with prewarmed (37°C) imaging media (Live Cell Imaging Solution, henceforth ‘imaging media’ (Thermo Fisher Scientific A14291DJ)) supplemented with 1% PSF 15 min prior to measurements.

For the blebbistatin treatment, cells were incubated with 20 μM blebbistatin (Sigma-Aldrich, B0560-1MG) for 15 min prior to measurement. For visualisation of membrane tethers, fibroblasts were stained with CellMask Orange Plasma Membrane Stain (ThermoFisher Scientific, USA, C10045-100 μl). The dye was added to the imaging media in the culture dish at 1:1000 dilution 15 min prior to the tether pulls.

### *Xenopus laevis* retinal ganglion cell culture

All animal experiments were conducted in compliance with the Ethical Review Committee of the University of Cambridge and Home office Guidelines.

*Xenopus* embryos were derived from *in vitro* fertilization of eggs kindly provided by the Gurden Institute Frog Facility (University of Cambridge, UK). For fluorescent labelling of *Xenopus* neurons, 5 nL of 0.2% w/v tetramethylrhodamine-labeled dextran (Thermo Fisher, D3308) diluted in nuclease-free water was injected in each dorsal blastomere of a 4-cell stage *Xenopus* embryo. Injections were performed as previously described^35^.

Embryos were grown at 14-16°C with daily cleaning in 0.1X Modified Barth’s Saline (10X MBS: 88mM NaCL, 2.4mM NaHCO3, 1mM HEPES, 0.82mM MgSO4, 0.33mM Ca(NO3)2, 0.41 mM CaCl2, pH 7.6).

All dissections took place at stage 35/36 (stages according to Nieuwkoop and Faber^36^). In the case of dextran-injected embryos, embryos were screened using a fluorescence stereomicroscope prior to dissection to ensure the presence of dextran in the eye primordia (note that dextran-injected embryos are morphologically normal). Embryos were anesthetized prior to dissection in 20% tricaine methanesulfonate solution (MS222, pH 7.6-7.8, with 1x PSF) and subsequently immobilized in a Sylgard®184-lined petri dish using a bent 0.2 mm insect bin (Austerlitz). Eye primordia were carefully explanted using a 0.15 mm insect pin (Austerlitz) held in a pin holder. Explants were immediately transferred to 60% L15 media (Sigma-Aldrich, L4386, pH 7.6-7.8, with 1x PSF, henceforth referred to as ‘Xenopus culture media’).

Following dissection, explants were placed with the lens facing up onto PAA gels or glass bottom culture dishes. The hydrogels were prepared as described below.

Hydrogels and glass dishes were coated overnight (hydrogels) or for 30 min (glass) with 0.01 mg/ml PDL (Sigma-Aldrich, P6407, diluted in PBS) followed by 2 h in 5 μg/mL laminin (Sigma-Aldrich, L2020, diluted in PBS). Following explant plating, dishes were allowed to sit for 1-2 h on the benchtop to allow explants to adhere, and then were moved to a 20°C incubator overnight.

### Immunostainings

Cells were washed twice with prewarmed (37°C) PBS. Afterwards, they were fixed and permeabilized with 4% PFA (Thermo Fisher 28908) and 0.2% tritonX (Sigma-Aldrich T8787) in PBS for 10 min at room temperature (RT). The cells were washed once with PBS at RT. Cells were incubated with DAPI (1.7 μg/ml Sigma-Aldrich D9542) and Phalloidin (Life Technologies A12379, diluted 1:300) in 1% Bovine Serum Albumin (Sigma-Aldrich A3294) in PBS for 1 h at RT. In the end, the dishes were washed twice with PBS. Cells were imaged with a Leica DMi8 epifluorescence setup with a ×63 oil objective (NA 1.4, Leica).

### Optical measurement

OT measurements took place on a custom-built inverted microscope^37^. Briefly, a 5 W 1064 nm ytterbium fibre laser (YLM-5-LP, IPG Laser, Germany) was used at a high power (> 1 W) to minimize intensity fluctuations. Excess power was redirected to a beam dump and the power at the optical trap itself was 600 mW. The trap was generated by overfilling the back aperture of an NA 1.2 UPlanSApo water immersion objective (Olympus, Japan) with the laser beam.

To enable controlled positioning of samples, the sample stage was mounted on top of a precision piezoelectric stage (P-517.43 and E-710.3, Physik Instrumente, Germany) with a positioning resolution of 1 nm. The sample stage was illuminated by a white fibre light source (DC-950 Fiber-Lite, Edmund Optics, USA). White light entered the objective and then passed through a dichroic mirror (DM) to a 550 nm short pass DM (DMSP550, Thorlabs, USA). This passed light below 550 nm to the 532 nm notch DM. This mirror coupled the fluorescence laser into the microscope, and in the process blocked the portion of incoming white light ranging from 515 to 540 nm. Light that passed through the notch DM was focused onto a CMOS camera (MC1362, Mikrotron, Germany), which was used for tracking particles in the optical trap. Light above 550 nm was passed to the fluorescence imaging camera (optiMOS Scientific CMOS, Teledyne Imaging, USA), which was used to find beads to trap, locate cells, and record fluorescence images of tethers.

Forces were measured with the OT by modelling a particle in the trap potential as a Hookean mass and spring system. By tracking the position of a trapped particle and multiplying this by a calibrated trap stiffness constant, the force experienced by the particle was determined. Trap calibration was performed using the power spectrum method^38^. Postprocessing of the data was done using a custom script in Python 3.6, which is available on GitHub https://github.com/mchughj33/mem-tension.

### OT bead functionalisation

As reported previously^39,40^, beads with 2 μm diameter (Kisker PC-S-2.0) were functionalized with Concanavalin A (ConA, from Canavalia, Sigma-Aldrich, C2272-10MG) as follows.

10 μL bead solution were added into 100 μL of 2 mg/mL ConA in a tube (low-protein binding) and incubated for 30 min at 4°C. Beads were washed twice via centrifugation in 100μL of 5mg/mL bovine serum albumin (BSA, Sigma-Aldrich, A3294) in 200mM Tris (Sigma, GE17-1321-01) in ddH2O. All centrifugation steps were conducted at 4°C, 10,000 rcf. After the final wash, 1 ml imaging media supplemented with 1% PSF was added and gently mixed by pipetting.

### Membrane tether pulling experiments

15 min prior to each fibroblast measurement, the culture medium was exchanged with pre-heated (37°C) imaging medium supplemented with 1% PSF. For Xenopus experiments, Xenopus medium was exchanged with RT imaging medium ~15 minutes prior to imaging.

In each OT experiment, 2 μl functionalised beads (see above) were added to the culture media and a bead captured in the trap. To reduce variations in the contact area between bead and cell^16^, all relevant measurement parameters were tightly controlled. Beads of one size (2.16 μm) were used, ConA coated beads were prepared freshly for each experiment under temperature-controlled conditions, and the beads were attached by bringing them into contact with the cells for the same amount of time (5 seconds, see below).

During the entire process, the position of the bead relative to the laser trap was recorded in order to measure the force produced by the tether pull. Before attaching the bead to the cell, a short recording of the Brownian motion of the trapped bead was taken. The mean force from each recording was used to define the point of zero force in the subsequent analysis of each respective pull. Afterwards, the bead was carefully brought into contact with the cell membrane. The image of the bead was monitored for a change in the interference pattern around the bead, indicating contact between the bead and membrane. At this point the beads was held stationary for 5 s. Then it was moved approximately 0.25 μm away from the cell while monitoring the particle tracking system to see if the mean displacement of the bead from the trap centre was reduced, which signified that the bead was attached to the membrane.

Beads were then moved away from the cell at 0.1 μm s in the case of 3T3 fibroblasts and 0.5 μm s^-1^ in the case of neurons, leading to the formation of a membrane tether between the cell and the bead. After reaching the end of the pull, 10 μm for 3T3 fibroblasts and 20 μm for neurons, the bead was held stationary for 30 s. The protocol for pulling tethers from neurons was based on previous work^41^. Due to high rate of tether loss during preliminary pulls with fibroblasts, to the pulling velocity was reduced, leading to a subsequent decrease in the pull distance to avoid reducing experimental throughput. Pontes et al. showed that varying the velocities for tether pulls in 3T3 fibroblasts within the range of 0.07 μm/s to 1μm/s, did not affect the force results^42^. We therefore do not expect this velocity change to affect our observations. For each pull, the peak force was defined as the maximum force value reached, and the steady state force was determined as the mean of the 30 s period at the end of the pull during which the bead was held stationary. Analysis was performed using custom analysis scripts written in python 3.6.

### Hydrogels for fibroblasts

Deformable hydroxy-polyacryamide substrates for fibroblast cultures were prepared as described previously^24,43^. Briefly, for each gel a top coverslip and a glass-bottomed culture dishe (MatTek P35G-0.20-C and Ibidi 81158 were used for OT and TFM, respectively) were prepared. Both coverslips were first washed in 70% ethanol and distilled water, and then dried. Bottom coverslips were wiped with a cotton bud soaked in 1M NaOH, dried, and then incubated in (3-aminopropyl)trimethoxysilane (APTMS, Sigma Aldrich, 281778) for 3 min. APTMS was washed off thoroughly in ddH20 and coverslips were then incubated in a 0.5% solution of glutaraldehyde (Sigma Aldrich, G6257) for 30 min and then rinsed 3x with distilled water. Top coverslips were treated for 10 min with Rain-X (Shell Car Care International). Acrylamide (AA, Sigma Aldrich, A4058) stock solution was made by mixing 500 μL of 40% Acrylamide + 65 μL of 100% Hydroxyacrylamide (Sigma Aldrich, 697931). Afterwards the gel premix was prepared by mixing 500 μL Acrylamide stock solution with 250 μL 2% Bisacrylamide (BA, Fisher Scientific, BP1404-250). The premix was diluted in PBS to achieve the different shear moduli according to the **table 1.** For TFM gels, 10 μL of PBS was replaced by 10 μl bead solution (FluoSpheres™ Carboxylate-Modified Microspheres, 0.2 μm, dark red fluorescent)) and the gel mix was sonicated for 30 s to separate the beads.

**Table 1:** Hydroxy-polyacrylamide gel composition used for fibroblasts

| Volume of gel | Volume of PBS | Shear modulus (G') |
| --- | --- | --- |
| premix |  |  |
| 53 μL | 447 μL | 0.1 kPa |
| 75 μL | 425 μL | 1 kPa |
| 150 μL | 350 μL | 10 kPa |

Freshly-made premixes were de-gassed under vacuum for 4-6 min. To each 500 μL of final gel solution, 1.5 μL 0.3% (v/v) N, N, N’, N’-tetramethylethylenediamine (TEMED, Thermo Fisher, 15524–010) and 5 μL 0.1% (w/v) ammonium persulfate (APS, Sigma, 215589) solution were added and the premix was gently mixed. 14 μL or 10 μL of this mixture were pipetted on the bottom coverslip and 18 mm or 16 mm top coverslips (for 14 μL and 10 μL volume, respectively) were placed on top (with RainX-treated side facing down). The gels were soaked in PBS for at least 20 min before the top coverslips were removed. Afterwards, they were washed twice with PBS and then treated with ultraviolet light for 30 min. Hydrogels were coated as described above.

### Hydrogels for neurons

Deformable polyacrylamide gel substrates were prepared as described previously^44,45^ for neuron culture. The coverslips were prepared as described in the hydroxy-acryamide gel protocol using MatTek dishes as the bottom coverslip. The gel mix was prepared according to **table 2** to achieve the different shear moduli. For TFM gels, the PBS volume was reduced by 10 μL and replaced by 10 μL bead solution and the gel mix was sonicated for 30 s to separate the beads.

**Table 2:** Polyacryamide gel composition used for neurons

| Volume of AA | Volume of BA | Volume of PBS | Shear modulus (G') |
| --- | --- | --- | --- |
| 63 μL | 10 μL | 427 μL | 0.1 kPa |
| 63 μL | 17.5 μL | 419.5 μL | 0.3 kPa |
| 94 μL | 15 μL | 391 μL | 1 kPa |

Freshly-made gel mixes were briefly de-gassed under vacuum for 4-6 min. To each 500 μL of final gel solution, 1.5 μL 0.3% (v/v) TEMED and 5 μL 0.1% (w/v) APS solution were added and gently mixed. 7 μL, 14 μL of this mixture were pipetted onto the bottom coverslip and 18 mm top coverslips were placed on top (with RainX treated side facing down) for OT experiments and TFM experiments, respectively. The gels were soaked in PBS for at least 20 min before the top coverslips were removed. The gels were subsequently functionalized via a 4h treatment with hydrazine hydrate (Sigma Aldrich, 225819), followed by 1 h in 5% acetic acid (Fisher Scientific, 10171460), and then washed thoroughly in PBS. Afterwards, they were treated with ultraviolet light for 30 min. Hydrogels were coated as described above.

### TFM

24 h after plating, fibroblasts were imaged with an inverted microscope (Leica DMi8) at 37°C and 5% CO_2_, equipped with a digital sCMOS camera (ORCA-Flash4.0, Hamamatsu Photonics), an EL6000 illuminator (Leica) and a ×63 oil objective (NA = 1.4, Leica). Images were acquired with Leica LAS X software. Fluorescence images of beads and wide-field images of cells were taken every 2 min for 10 minutes. After image acquisition, the culture media was exchanged with Trypsin-EDTA (Gibco) to detach cells from the gel. Reference images of fluorescent beads were taken 15 min after trypsinization.

16 - 24 h after plating, *Xenopus* eye primordia explants were imaged using an inverted Nikon microscope (Ti-E at room temperature) equipped with an sCMOS camera (Prime BSI Scientific, Photometrics), a CoolLED pE-4000, and ×60 oil objective (NA = 1.4, Nikon). Fluorescence images of beads and wide-field images of cells were taken every 5 min for 1 h. After image acquisition, the culture media was exchanged with Trypsin-EDTA (Gibco) or 20% sodium dodecyl sulfate solution (Severn Biotech Ltd., 2 0-4002-01) to detach the explants from the gel. Reference images of fluorescent beads were taken 15 min after trypsinization or immediately after sodium dodecyl sulfate treatment.

Traction stress maps were calculated for each frame. Traction stresses were averaged over time (12 frames for neurons*, 5* frames for fibroblasts) for each cell. Postprocessing of the data and statistical analyses were done with a custom script in Python 3.8 and is available on GitHub https://github.com/rg314/pytraction.

### Statistics

OT data are shown as violin plots, TFM data as box plots. Violin shapes are kernel density estimations, boxes indicate the range of data that falls between the first and third quartile, whiskers show the range of data within 1.5 times the interquartile range. Density estimator width is based on the number of samples at a given value. Median values for each dataset are marked by a white circle. Lines between violins indicate significant differences in the samples. The boxplot diagrams were generated using python seaborn. The borders of the box indicate the quartiles of the dataset, the median is indicated by a line within the box, and the whiskers include 1.5*interquartile range. Points that were determined to be outliers were plotted individually. Kruskal-Wallis one-way analysis of variance (ANOVA) tests followed by a Tukey posthoc test were used for statistical analysis when comparing multiple groups. When comparing two groups, a Mann-Whitney test was used.

## References

1. Raucher, D. & Sheetz, M. P. Cell Spreading and Lamellipodial Extension Rate Is Regulated by Membrane Tension. J. Cell Biol. 148, 127–136 (2000).

2. Nussenzveig, H. M. Cell membrane biophysics with optical tweezers. Eur. Biophys. J. 47, 499–514 (2018).

3. Gauthier, N. C., Fardin, M. A., Roca-Cusachs, P. & Sheetz, M. P. Temporary increase in plasma membrane tension coordinates the activation of exocytosis and contraction during cell spreading. Proc. Natl. Acad. Sci. U. S. A. 108, 14467–14472 (2011).

4. Keren, K. et al. Mechanism of shape determination in motile cells. Nature 453, 475–480 (2008).

5. De Belly, H. et al. Membrane Tension Gates ERK-Mediated Regulation of Pluripotent Cell Fate. Cell Stem Cell 28, 273–284.e6 (2021).

6. Dai, J. & Sheetz, M. P. Regulation of endocytosis, exocytosis, and shape by membrane tension. Cold Spring Harb. Symp. Quant. Biol. 60, 567–571 (1995).

7. Sheetz, M. P. Cell control by membrane–cytoskeleton adhesion. Nat. Rev. Mol. Cell Biol. 2, 392–396 (2001).

8. Pontes, B., Monzo, P. & Gauthier, N. C. Membrane tension: A challenging but universal physical parameter in cell biology. Semin. Cell Dev. Biol. 71, 30–41 (2017).

9. Pontes, B. et al. Membrane tension controls adhesion positioning at the leading edge of cells. J. Cell Biol. 216, 2959–2977 (2017).

10. Salbreux, G., Charras, G. & Paluch, E. Actin cortex mechanics and cellular morphogenesis. Trends Cell Biol. 22, 536–545 (2012).

11. Clark, A. G., Wartlick, O., Salbreux, G. & Paluch, E. K. Stresses at the Cell Surface during Animal Cell Morphogenesis. Curr. Biol. 24, R484–R494 (2014).

12. Califano, J. P. & Reinhart-King, C. A. Substrate Stiffness and Cell Area Predict Cellular Traction Stresses in Single Cells and Cells in Contact. Cell. Mol. Bioeng. 3, 68–75 (2010).

13. Sabass, B., Gardel, M. L., Waterman, C. M. & Schwarz, U. S. High Resolution Traction Force Microscopy Based on Experimental and Computational Advances. Biophys. J. 94, 207–220 (2008).

14. Sheetz, M. P. Cell control by membrane–cytoskeleton adhesion. Nat. Rev. Mol. Cell Biol. 2, 392–396 (2001).

15. Datar, A., Bornschlögl, T., Bassereau, P., Prost, J. & Pullarkat, P. A. Dynamics of membrane tethers reveal novel aspects of cytoskeleton-membrane interactions in axons. Biophys. J. 108, 489–497 (2015).

16. Koster, G., Cacciuto, A., Derényi, I., Frenkel, D. & Dogterom, M. Force barriers for membrane tube formation. Phys. Rev. Lett. 94, 16–19 (2005).

17. Sens, P. & Plastino, J. Membrane tension and cytoskeleton organization in cell motility. J. Phys. Condens. Matter 27, (2015).

18. Gauthier, N. C., Masters, T. A. & Sheetz, M. P. Mechanical feedback between membrane tension and dynamics. Trends Cell Biol. 22, 527–535 (2012).

19. Paraschiv, A. et al. Influence of membrane-cortex linkers on the extrusion of membrane tubes. Biophys. J. 120, 598–606 (2021).

20. Xu, K., Zhong, G. & Zhuang, X. Actin, spectrin, and associated proteins form a periodic cytoskeletal structure in axons. Science 339, 452–456 (2013).

21. Wells, R. G. Tissue Mechanics and Fibrosis. Biochim. Biophys. Acta 1832, 884–890 (2013).

22. Franze, K., Janmey, P. A. & Guck, J. Mechanics in Neuronal Development and Repair. Annu. Rev. Biomed. Eng. 15, 227–251 (2013).

23. Koch, D., Rosoff, W. J., Jiang, J., Geller, H. M. & Urbach, J. S. Strength in the Periphery: Growth Cone Biomechanics and Substrate Rigidity Response in Peripheral and Central Nervous System Neurons. Biophys. J. 102, 452–460 (2012).

24. Rheinlaender, J. et al. Cortical cell stiffness is independent of substrate mechanics. Nat. Mater. 19, 1019–1025 (2020).

25. Schwarz, U. S. & Soiné, J. R. D. Traction force microscopy on soft elastic substrates: A guide to recent computational advances. Biochim. Biophys. Acta BBA - Mol. Cell Res. 1853, 3095–3104 (2015).

26. Huang, Y. et al. Traction force microscopy with optimized regularization and automated Bayesian parameter selection for comparing cells. Sci. Rep. 9, 539 (2019).

27. Franze, K. Integrating Chemistry and Mechanics: The Forces Driving Axon Growth. Annu. Rev. Cell Dev. Biol. 36, 61–83 (2020).

28. Lieber, A. D., Yehudai-Resheff, S., Barnhart, E. L., Theriot, J. A. & Keren, K. Membrane Tension in Rapidly Moving Cells Is Determined by Cytoskeletal Forces. Curr. Biol. 23, 1409–1417 (2013).

29. Mandal, K. et al. Soft hyaluronic gel promotes cell spreading, stress fibers, focal adhesion, membrane tension by phosphoinositide signaling, not traction force. ACS Nano 13, 203–214 (2019).

30. Thottacherry, J. J. et al. Mechanochemical feedback control of dynamin independent endocytosis modulates membrane tension in adherent cells. Nat. Commun. 9, 1–14 (2018).

31. Vignaud, T. et al. Stress fibres are embedded in a contractile cortical network. Nat. Mater. 1–11 (2020) doi:10.1038/s41563-020-00825-z.

32. Shi, Z., Graber, Z. T., Baumgart, T., Stone, H. A. & Cohen, A. E. Cell membranes resist flow. Cell 175, 1769–1779.e13 (2018).

33. Ellefsen, K. L. et al. Myosin-II mediated traction forces evoke localized Piezo1-dependent Ca 2+ flickers. Commun. Biol. 2, 1–13 (2019).

34. Walcott, S. & Sun, S. X. A mechanical model of actin stress fiber formation and substrate elasticity sensing in adherent cells. Proc. Natl. Acad. Sci. 107, 7757–7762 (2010).

35. Leung, K.-M. et al. Asymmetrical ß-actin mRNA translation in growth cones mediates attractive turning to netrin-1. Nat. Neurosci. 9, 1247–1256 (2006).

36. Nieuwkoop, P. D. & Faber, J. Normal table of xenopus laevis (daudin). (Routledge, 1994).

37. Mc Hugh, J., Andresen, K. & Keyser, U. F. Cation dependent electroosmotic flow in glass nanopores. Appl. Phys. Lett. 115, 113702 (2019).

38. Gittes, F. & Schmidt, C. F. Chapter 8 signals and noise in micromechanical measurements. in Methods in cell biology. vol. 55 129–156 (Academic Press, 1997).

39. Buer, C. S., Weathers, P. J. & Swartzlander, G. A. Changes in Hechtian strands in cold-hardened cells measured by optical microsurgery. Plant Physiol. 122, 1365–1377 (2000).

40. Kucik, D. F., Kuo, S. C., Elson, E. L. & Sheetz, M. P. Preferential attachment of membrane glycoproteins to the cytoskeleton at the leading edge of lamella. J. Cell Biol. 114, 1029–1036 (1991).

41. Hochmuth, R. M., Shao, J. Y., Dai, J. & Sheetz, M. P. Deformation and flow of membrane into tethers extracted from neuronal growth cones. Biophys. J. 70, 358–369 (1996).

42. Pontes, B. et al. Cell cytoskeleton and tether extraction. Biophys. J. 101, 43–52 (2011).

43. Moshayedi, P. et al. The relationship between glial cell mechanosensitivity and foreign body reactions in the central nervous system. Biomaterials 35, 3919–3925 (2014).

44. Koser, D. E. et al. Mechanosensing is critical for axon growth in the developing brain. Nat. Neurosci. 19, 1592–1598 (2016).

45. Moshayedi, P. et al. Mechanosensitivity of astrocytes on optimized polyacrylamide gels analyzed by quantitative morphometry. J. Phys. Condens. Matter 22, (2010).

