## Supplementary figures for "Effective cell membrane tension is independent of polyacrylamide substrate stiffness"

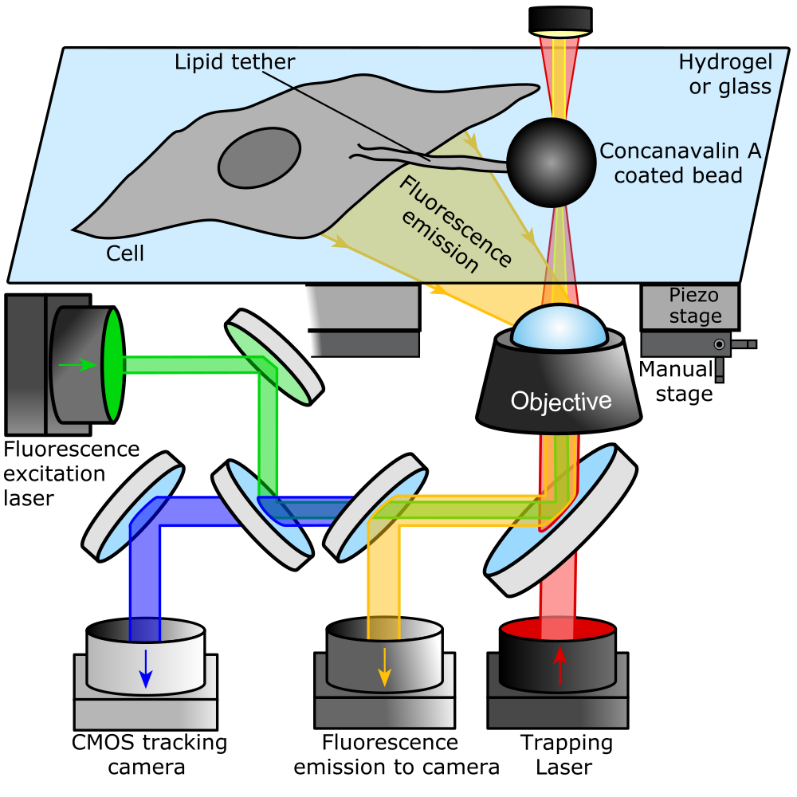

**Figure S1:** **Schematic of the optical tweezers setup.** The optical trap (OT) trapped a bead coated with concanavalin A. The bead was positioned close to a cell’s membrane using a manual microscope stage. After approximately 5 seconds, the bead was moved away from the cell using a piezo stage. The deflection of the bead due to a membrane tether was recorded with a CMOS camera. When the pull was complete, fluorescence microscopy was used to visualise the tether (Figs. 2a, e).

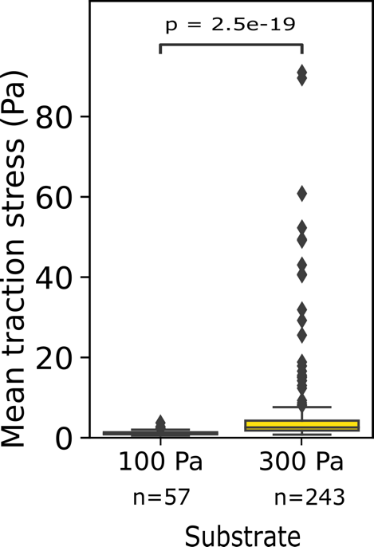

**Figure S2:** **Traction forces of neuronal growth cones.** The cells exerted higher traction forces on stiff hydrogels compared to softer hydrogels (two-tailed Mann-Whitney test).

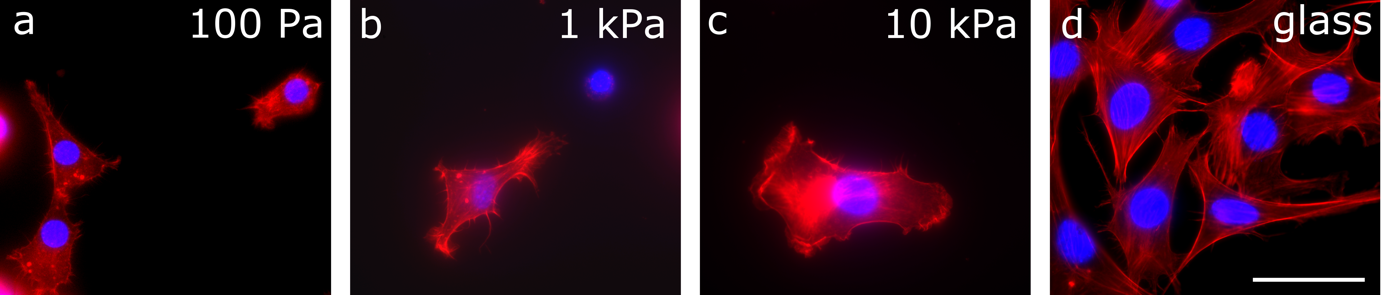

**Figure S3:** **Immunostainings of fibroblasts on different substrates**. Fibroblasts grown on hydrogels and glass, stained for actin (red) and nuclei (blue). Note the drastic change in morphology between hydrogel and glass substrates. Cells on glass exhibited pronounced stress fibres. Scale bar: 50µm.
